# NGSEP 4: Efficient and Accurate Identification of Orthogroups and Whole-Genome Alignment

**DOI:** 10.1101/2022.01.27.478091

**Authors:** Daniel Tello, Laura Natalia Gonzalez-Garcia, Jorge Gomez, Juan Camilo Zuluaga-Monares, Rogelio Garcia, Ricardo Angel, Daniel Mahecha, Erick Duarte, Maria del Rosario Leon, Fernando Reyes, Camilo Escobar-Velásquez, Mario Linares-Vásquez, Nicolas Cardozo, Jorge Duitama

## Abstract

Whole-genome alignment allows researchers to understand the genomic structure and variations among the genomes. Approaches based on direct pairwise comparisons of DNA sequences require large computational capacities. As a consequence, pipelines combining tools for orthologous gene identification and synteny have been developed. In this manuscript, we present the latest functionalities implemented in NGSEP 4, to identify orthogroups and perform whole genome alignments. NGSEP implements functionalities for identification of clusters of homologus genes, synteny analysis and whole genome alignment, and visualization. Our results showed that the NGSEP algorithm for ortholog identification has competitive accuracy and better efficiency in comparison to commonly used tools. The implementation also includes a visualization of the whole genome alignment based on synteny of the orthogroups that were identified, and a reconstruction of the pangenome based on frequencies of the orthogroups among the genomes. Finally, our software includes a new graphical user interface. We expect that these new developments will be very useful for several studies in evolutionary biology and population genomics.

## INTRODUCTION

Whole-genome alignment (WGA) is the process of identifying high-quality multiple local alignments within a collection of assembled genomes. It allows a direct comparison of two or more complete genomes within or between species. This comparison permits researchers not only to find regions of collinearity between the genomes, but also to find different alleles within a population (Minkin and Medvedev 2020). The WGA problem is also closely related to finding homologous genes, to reconstruct the synteny, and to represent pan-genomes. To solve this problem, two main strategies have been developed: one based on pairwise local alignments and another based on large collinear blocks. The first strategy finds all local similarities between pairs of genomes and then combines these pairwise alignments into multiple alignments. The second one identifies sets of large collinear homologous segments among the analyzed genomes, and then these blocks are independently aligned (Dewey and Pachter 2006). Several tools have been released to perform different parts of the genome alignment process (Supplementary Table 1). However, none of these tools integrate orthologs identification, synteny, and alignment in an efficient and easy to use software solution.

Classical solutions to align large DNA sequences include MUMmer (Delcher et al. 1999) and Mauve (Darling et al. 2004). The core of both algorithms is the identification of maximal unique matches (MUMs) within the genomes that are being compared. Whereas MUMmer implements a suffix tree, Mauve builds a table with the location of all k-mers within the sequences. Both approaches are inefficient to compare mammalian genomes, especially in the amount of random access memory (RAM) required to build the structures needed to calculate the MUMs. In a more recent tool called Satsuma, sequences are transformed to treat them as audio signals and similarities are identified by cross-correlation using the fast Fourier transform (Grabherr et al. 2010); although this approach seeks to overcome issues related to biases produced by seeded alignments and by repetitive regions, this solution is inefficient to compare large genomes. The availability of a growing number of complex genomes has motivated the development of novel genome comparison tools in recent years. As an example, closely-related mammal genomes were successfully aligned using SibeliaZ-LCB, which implements compacted de Bruijn graphs to find collinear blocks, reducing the consumption of computational resources and time (Minkin and Medvedev 2020). Although current solutions are more efficient than MUMmer and Mauve, one general issue with sequence-based genome alignment tools is that it is not easy for users to identify orthology of important genomic elements such as genes. Regarding visualization, only Mauve provides a solution for visualization of their Locally Collinear Blocks (LCBs) of synteny. Tools such as the Artemis Comparison Tool (Carver et al. 2005) and Symap (Soderlund et al. 2011) were built to visualize direct blast searches and the results of MUMmer respectively.

An alternative solution to align and compare whole-genomes relies on the identification of groups of homologous genes among the genomes (called orthogroups). These orthogroups are aligned using information of physical adjacency. Two widely used tools to determine orthologous groups are Orthofinder (Emms and Kelly 2019) and SonicParanoid (Cosentino and Iwasaki 2019). Both tools are based on reciprocal best hits, obtained using heuristic pairwise aligners, such as BLAST. Orthofinder implements a normalization of BLAST scores and then performs a Markov Clustering (Emms and Kelly 2015). Using ortholog groups, Orthofinder then infers gene trees and tries to identify the rooted species-tree, allowing the algorithm to split ortholog groups according to gene duplication events (Emms and Kelly 2019). In contrast, SonicParanoid replaces BLAST with MMSeq2 to speed-up the alignment and clustering steps and implements a graph-based algorithm to identify ortholog groups (Cosentino and Iwasaki 2019). Although both software tools perform well in identifying orthogroups, their input is the proteome derived from the genomes which results in a lack of information about adjacency and synteny.

This manuscript describes the new functionalities implemented in NGSEP to perform alignments of complete genomes and identification of ortholog clusters. In summary, the new functionality performs the following tasks: 1) Clustering of genes in orthogroups; 2) whole genome alignment; and 3) interactive web-based visualization of the alignment. The reliability and efficiency of our software was tested against currently available open-source software for genome alignment.

## NEW FUNCTIONALITIES FOR COMPARATIVE GENOMICS

The development of the genomes aligner was inspired in recent works in which different tools need to be integrated to perform comparative genomics between genome assemblies (See for instance (Moghaddam et al. 2021)). The input of this process is either a set of annotated genomes (FASTA files with gene annotations in GFF3 format), a set of DNA coding sequences (CDS) or a set of aminoacid sequences. Proteomes for each genome are extracted and homology relationships (orthologs and paralogs) are efficiently predicted by indexing of aminoacid sequences by k-mers and calculation of percentages of shared k-mers. Then, genes are clustered in orthogroups based on the topology of the graph induced by the predicted relationships. Finally, a presence/absence matrix is derived from these orthogroups and the gene families are classified as soft/exact core and accessory.

If genome assemblies are provided as input, synteny relationships are identified for each pair of genomes implementing an adapted version of the HalSynteny algorithm (Krasheninnikova et al. 2020). Homology relationships, classified orthogroups and synteny relationships are reported in a set of text files. Moreover, an interactive visualization of synteny blocks and ortholog relationships is provided as an HTML file. Figure 1 shows a schematic of the entire process. These algorithms are provided as functionalities of the Next Generation Sequencing Experience Platform (NGSEP) (Duitama et al. 2014) and they are available in the new graphical user interface and in the command line interface (commands CDNACatalogAligner and GenomesAligner). We describe the details of the implementation below.

**Figure 1.**
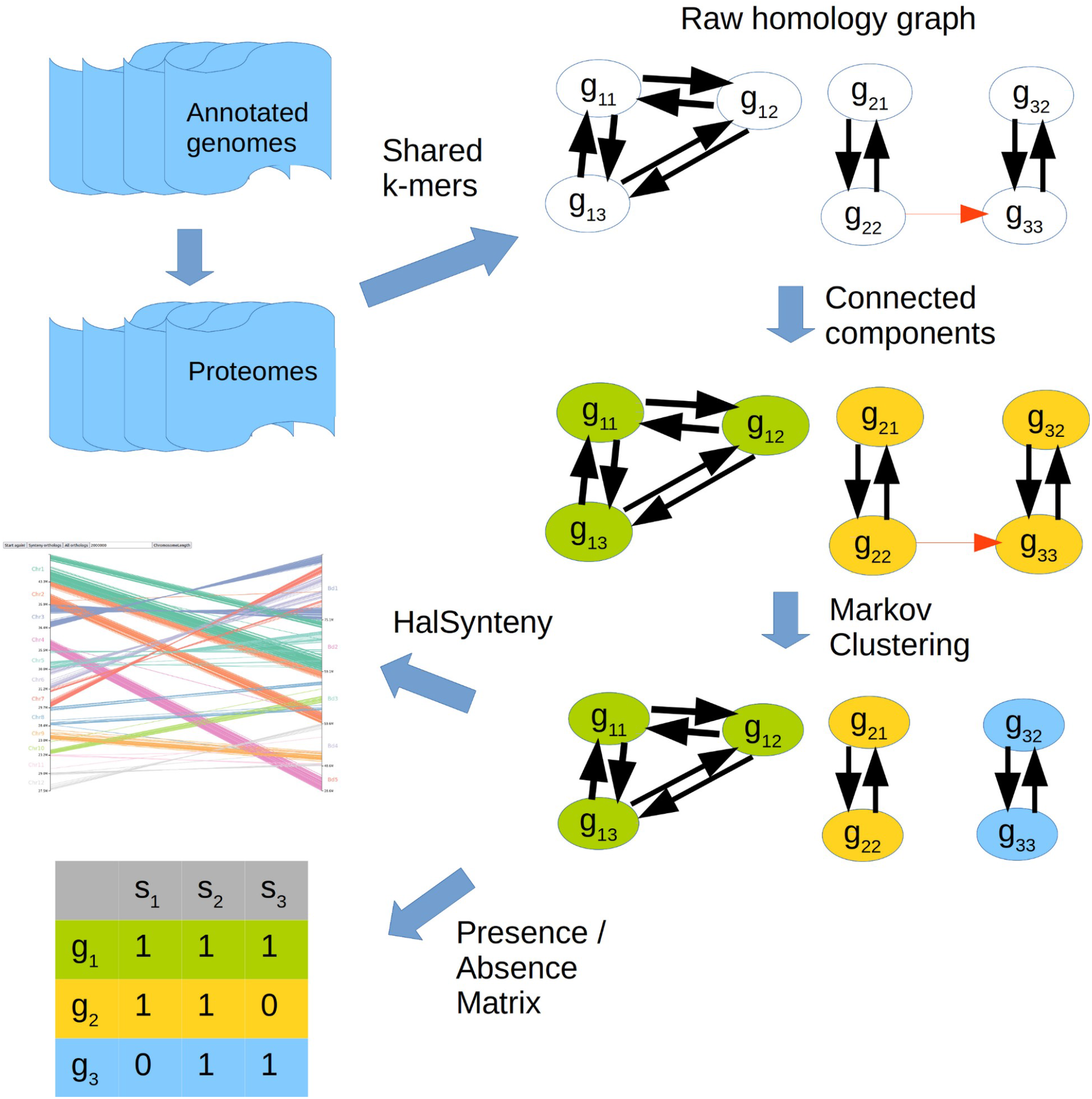
Schematic of the functionalities for comparative genomics implemented in NGSEP 4. Proteomes are either received directly as input or inferred from annotated genomes. A graph of raw candidate homolog relationships is built from the percentage of shared k-mers between each pair of genes. The red arrow corresponds to a false positive relationship. Clustering is performed from the graph running Markov Clustering on the connected components. If genomes are provided, the HalSynteny algorithm is used to identify synteny relationships between each pair of genomes. A presence/absence matrix is inferred from the homology clusters

### Identification of homology relationships and clustering in orthogroups

The orthogroups identification module takes as input a set of proteomes from different samples (usually species), either provided directly by the user, or reconstructed from annotated genome assemblies or CDS. For genome assemblies, the complete list of transcripts is recovered and for each gene, the longest transcript is extracted to get as much information as possible. In both cases, a homology unit is generated for each protein.

To identify both ortholog and paralog relationships, a table of k-mers is built storing for each k-mer pointers to the sequences containing such k-mer. Afterwards, k-mers from each amino acid sequence in a proteome are matched against the table of such proteome to calculate the percentage of shared k-mers between each pair of proteins within the proteome. Two proteins are considered paralogs if they share more than a user-specified percentage of k-mers. Likewise, orthologs are identified matching non overlapping k-mers of each query amino acid sequence of one proteome against the k-mers table of the subject proteome. The percentage of shared k-mers between query and subject proteins are calculated and the potential relationships are filtered using the same threshold applied for paralogs. Both paralog and ortholog relationships induce a directed graph with homology units (amino acid sequences) as vertices and relationships as edges. Prior to clustering, and based on the symmetry property of orthologs, the directed graph is converted into an undirected graph adding the missing edges with a default score of 0 to allow the clustering of both genes into the same orthogroup.

To identify orthogroups among a set of proteomes, breadth-first search is executed on the orthology unit graph to identify connected components. Looking for computational efficiency, the initial calculation of percentages of shared k-mers is an estimate from non-overlapping k-mers. Hence, in this step, percentages are recalculated using all k-mers for each pair of sequences clustered in the same connected component. This operation transforms the graph into a set of complete subgraphs. Edges with percentage of k-mers lower than the given threshold are removed and connected components are recalculated to refine the initial clustering obtained from raw predictions. Furthermore, the Markov Clustering (MCL) algorithm implemented in OrthoMCL (Li et al. 2003) was reimplemented in NGSEP and executed within each connected component to break incorrect clusters generated by false homology predictions. Each connected component is then passed through the MCL algorithm as an individual task returning a set of clusters. In brief, the MCL estimates the flow within the connected component. This flow is represented by random walks throughout the component where an agent can traverse the graph visiting edges according to transition probabilities. These probabilities are defined by the row normalization of the scores of ortholog relationships in the connected component. Subclustering of the component is performed via the inflation-expansion process. The inflation process raises each entry in the matrix to a power (two in this case), and normalizes the results to increase the probability of strong edges and decrease the probability of weak edges. The expansion process calculates the probabilities of the agent to move from a source node to a target node in n steps (8 in this case). When inflation and expansion are applied in alternance to the transition matrix, it will converge into a state where the resulting flows are clear, and each row indicates the cluster assigned to each node. The resulting separated clusters are considered the final orthogroups. Finally, genes without a cluster are added as private clusters.

In order to catalog orthogroups into core/accessory genomes, a presence/absence (PA) matrix is computed and frequencies of orthogroups within the genomes are calculated. Frequencies are then used to classify each orthogroup based on a threshold. A user defined threshold (90% by default) is applied to separate the core genome from the accessory genome.

### Genome alignment, synteny and visualization

The orthogroups obtained from the clustering analysis are used as input to perform alignment between all possible pairs of genomes. Given two genomes, NGSEP implements a modified version of the HalSynteny algorithm (Krasheninnikova et al. 2020) to identify syntenic relationships and perform genome alignment. In brief, this algorithm builds a graph with ortholog pairs as vertices. Given two vertices v1=<a1,b1> and v2=<a2,b2>, where a1, a2 are genes in the first genome and b1, b2 are genes in the second genome, a directed edge is added between v1 and v2 if a2 is located after a1 in the same chromosome, b1 and b2 are located in the same chromosome, the distance between a1 and a2 is lower than a given threshold, and the absolute distance between b1 and b2 is lower than the same threshold. Vertices are sorted (topologically) based on the coordinates in the first genome. Then, a synteny block is identified making a single traversal of the vertices, and calculating for each vertex the total length of the longest path that finishes in such vertex. The predecessor of each best path is also stored for each vertex. The vertex with the longest global path length is chosen as the last vertex of the synteny path and predecessors are used to reconstruct the path. Vertices of the chosen path are removed and the process is executed on the remaining subgraph until the chosen path has total length below a given threshold.

As main modifications to the algorithm described in (Krasheninnikova et al. 2020), adjacent paralogs probably generated by local duplication events are collapsed into a single “ local orthogroup”. This makes the algorithm more robust to recent duplication events. Also, large orthogroups in which the product between local orthogroups in genome 1 and the local orthogroups in genome 2 are larger than 100 are ignored. These cases correspond to large gene families spread over the genome, which are not likely to be informative for synteny analysis.

Finally, we implemented a web-enabled interactive visualizer for whole genome alignments based on the D3 framework (https://d3js.org/), which is one of the currently leading technologies for visualization of large datasets. The visualization shows the two genomes being compared as vertical columns parallel to each other, with their respective chromosomes labeled. Synteny blocks are visualized according to their relative direction. The visualizer allows the user to interact with it by selecting any chromosome on either genome and revealing only the synteny found within a selected region. We added buttons to allow visualization of individual ortholog relationships. Figure 2 shows the default visualization for the synteny between rice (*Oryza sativa*) and *Brachipodium distachion*, as reconstructed by the genomes aligner. The reconstruction for the genomes of human and chimpanzee can be visualized in the Supplementary Figure S1.

**Figure 2.**
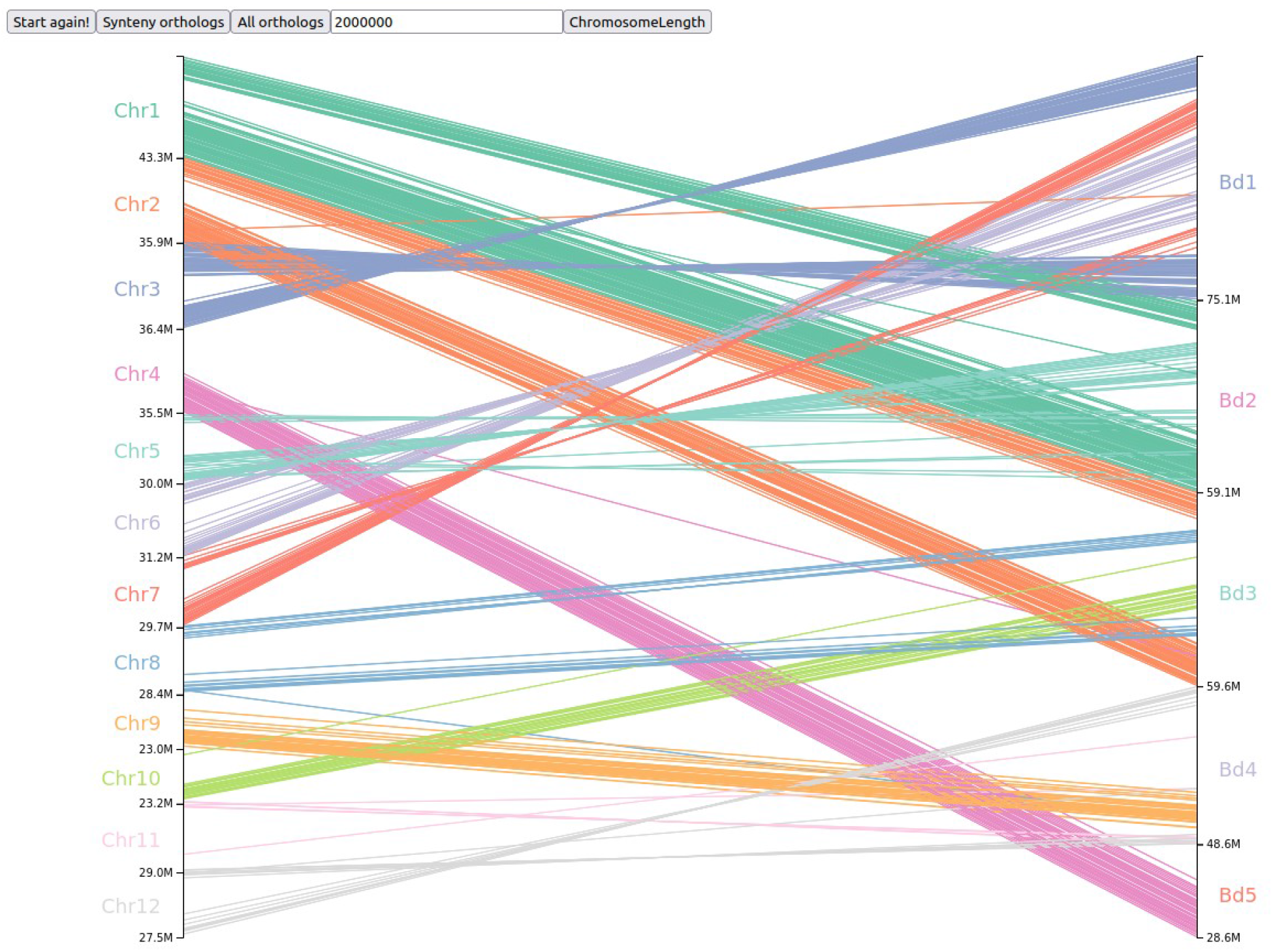
Comparison between the rice genome (*O. sativa*, left) and the genome of *Brachipodium distachion* (right), as generated by the genomes aligner of NGSEP. Synteny blocks are colored by rice chromosome

### Graphical user interface

We developed a new interface for NGSEP, based on JavaFX and completely independent from Eclipse, which was the technological framework of the previous user interface. When developing this new interface, we had two goals in mind: making the installation process simpler and independent from external software while also delivering a user friendly and easy to use interface with an updated new look. To achieve the goal of facilitating the installation of the application and its graphical component, we built this interface based on the JavaFX technology, which is an open-source software framework to implement user interfaces for Java applications that could be deployed either as web or desktop applications. With the use of JavaFX SDK, we were able to generate a package with a runnable file for each operative system, having the standard Java 11 JRE as the only prerequisite for successful installation.

We integrated in this new interface the features previously available in the Eclipse plugin, including a general file explorer, a custom file viewer with preview visualization mode for large gzip compressed Fasta and Fastq files as well as read alignment files in BAM format, and a progress bar and stop button to keep control of asynchronous processes. An important advantage of this new look and feel is that all the unnecessary buttons and generic functions of the Eclipse plugin are not present in this implementation, facilitating the general usage for researchers working with sequencing reads. The Supplementary Figure S2 shows a screenshot of this new interface.

## TESTING AND BENCHMARK EXPERIMENTS

### Parameter tuning guided by Orthobench

A set of 70 manually curated orthogroups available in the Orthobench dataset (Emms and Kelly 2020) was retrieved to tune parameters of our algorithm and perform comparisons against current software tools for identification of ortholog relationships (See methods for details). To explore the feasibility of ortholog clustering by percentage of shared k-mers, we calculated the distributions of shared k-mers between all pairs of sequences within orthogroups for different values of k (Supplementary Figure S3). As expected, the percentage of shared k-mers for highly divergent reference orthogroups is much lower than that of the slightly divergent orthogroups.

Based on the previous result, we tested the NGSEP clustering over a set of sequences belonging to 70 orthogroups from 12 species of Bilateria, varying the k-mer length from 5 to 9 and the percentage of shared k-mers from 1% to 10%. Recall, precision and F-score were calculated for each orthogroup (Figure 3A and Supplementary Table S2). The same analysis was performed filtering the dataset to include only mammalian species. In both cases, the best F-score was obtained for k=5 and 5% shared k-mers using the Markov clustering algorithm (blue triangle, Figure 3A). Unexpectedly, when the Markov clustering step is not used, the connected components algorithm performs better with k=9 and p=2% (Supplementary Table S2). The cause of this result is that the connected components algorithm tends to have a higher recall whilst having lower precision. Using a higher value of k and lower percentage of shared k-mers allows to target specific conserved regions (i.e. domains) within highly divergent sequences. The execution time for this dataset is lower than 0.1 seconds except on k-mer lengths below 6 and percentages below 2%, as shown in Figure 3b. This is due to the creation of large connected components, which has a longer execution time in the Markov clustering algorithm.

**Figure 3.**
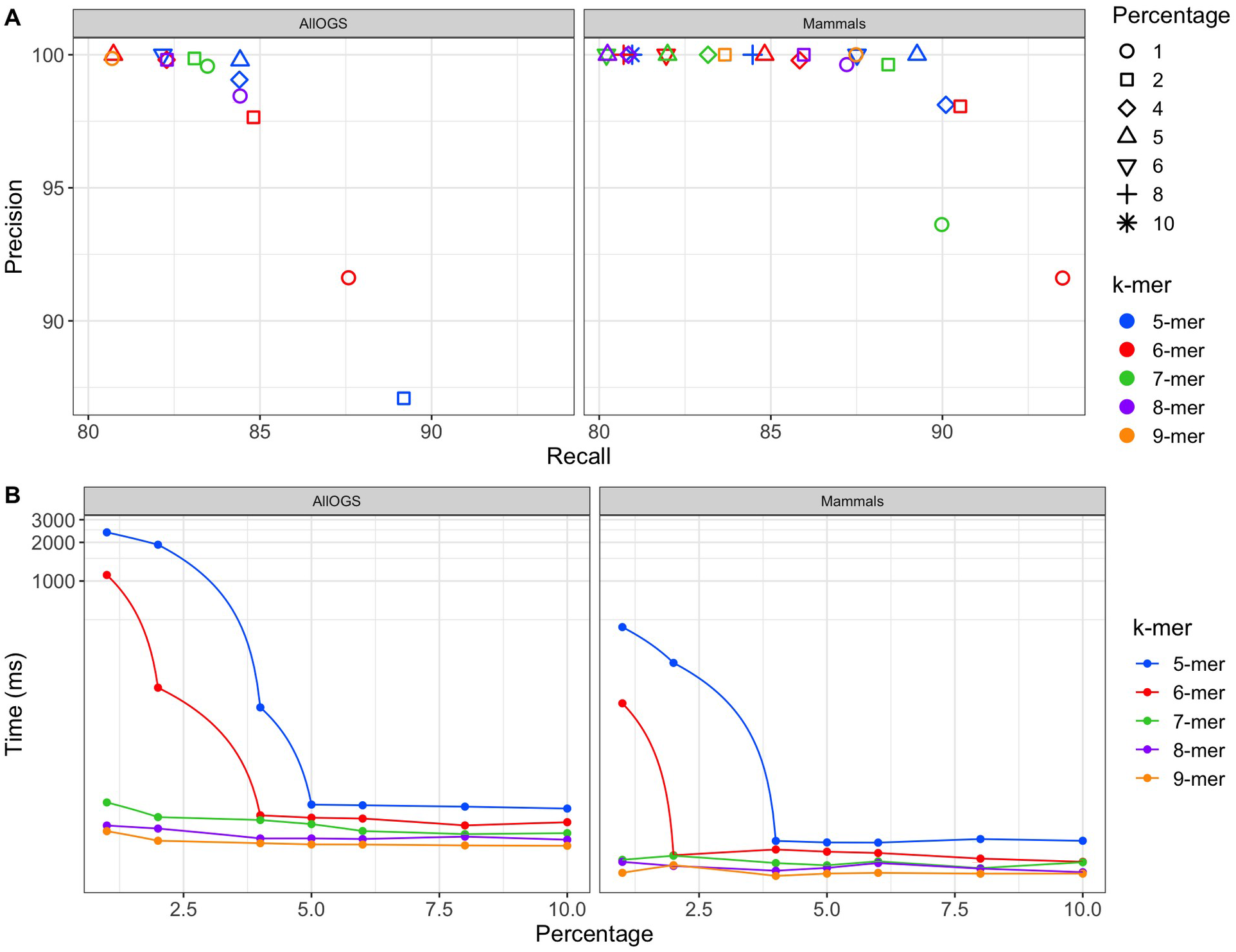
Markov clustering classification results using different combinations of parameters. A. Precision and Recall; B. Time consumption using 1 thread.

### Benchmarking orthologs

Using the tuned parameters, we also compared the implemented clustering approach against the results of two recently released software tools: OrthoFinder (version 1 and 2) (Emms and Kelly 2019) and SonicParanoid (Cosentino and Iwasaki 2019). These are the best performing tools to determine orthology groups according to the benchmarking service of Orthobench. Precision, recall, F-score, and runtimes for each tool are presented in Table 1. The best F-score was obtained with Orthofinder1 based on BLAST (92.7), followed by Orthofinder1 based on Diamond (92.4). NGSEP ranked third (91.5), overcoming SonicParanoid (86). The result of NGSEP was less than 1% lower from the best result, and its execution time was at least 10 times faster than Orthofinder and SonicParanoid.

**Table 1:**
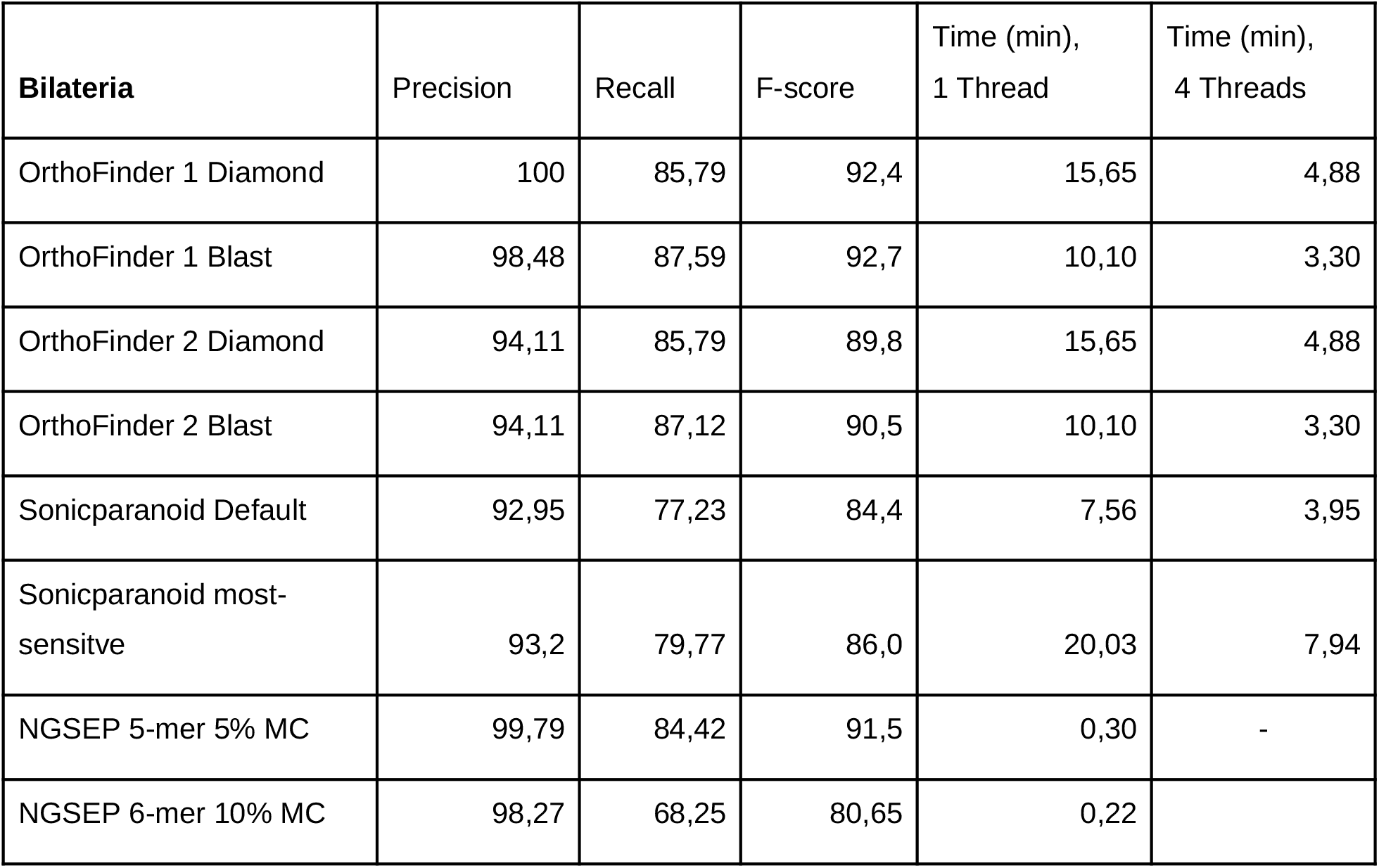
Benchmarking statistics of orthologs identification using the curated orthologs covering Bilateria from Orthobench.

The same analysis was performed over the mammals dataset, and the results for each tool are presented in Table 2. NGSEP obtained the best performance with an F-score of 94.3, being 100% precise and more sensitive than the other tools. This result is almost 10 points over the second best result that was obtained with Orthofinder using Diamond (F-score 85). Using this dataset, the F-score decreased because of the sensitivity using all tools, with the exception of NGSEP. Regarding computational efficiency NGSEP kept the best execution time (0.15 minutes or 9 seconds), which was more than 10 times faster than the second fastest tool (SonicParanoid with default parameters).

**Table 2:** Benchmarking statistics of orthologs identification only using the mammals orthologs sequences from Orthobench.

| **Mamalia** | Precision | Recall | F-score | Time (min),<br>1 Thread | Time (min),<br>4 Threads |
| --- | --- | --- | --- | --- | --- |
| OrthoFinder 1 Diamond | 100 | 73,86 | 85,0 | 4,83 | 1,84 |
| OrthoFinder 1 Blast | 100 | 73,5 | 84,7 | 4,16 | 1,45 |
| OrthoFinder 2 Diamond | 86,55 | 63,57 | 73,3 | 4,83 | 1,84 |
| OrthoFinder 2 Blast | 87,03 | 66,13 | 75,2 | 4,16 | 1,45 |
| Sonicparanoid Default | 86,45 | 62,48 | 72,5 | 2,04 | 0,86 |
| Sonicparanoid most-sensitive | 86,5 | 62,62 | 72,6 | 6,90 | 2,49 |
| NGSEP 5-mer 5% MC | 100 | 89,26 | 94,3 | 0,15 | - |
| NGSEP 6-mer 10% MC | 100 | 78,46 | 87,9 | 0,11 | - |

### Comparison of bacterial gene families

To evaluate the software usability building pangenomes, *Escherichia coli* was used as a model due to the availability of more than 15,000 complete genomes. Assemblies were retrieved from GenBank and pangenomic presence/absence matrices were built using subsamples of 10 to 100 genomes. The exact/soft core and accessory genomes were obtained based on the orthogroups identification and frequencies. The results of each partition fitted a normal distribution according to the Shapiro-Wilk normality test (p-value >0.05). As shown in Figure 4a, the exact core genome reduces from more than 3000 to less than 2000 orthogroups as the number of genomes increases, whereas the soft-core genome only reduces from around 3500 to around 3100 orthogroups when 100 genomes are included. Additionally, the Student’s t test showed a significant difference in the mean of orthogroups between the soft and exact approaches in all cases (p-value < 1.9e-15). Comparing the results with different amounts of genomes, we observed that in most of the cases over 50 genomes, there is no significant difference in the mean of orthogroups assigned to the soft-core genome, according to the Welch Two Sample test (p-value < 0.05, Supplementary Table S2). However, there is a significant difference comparing the exact core genome (p-value < 0.05 except in the comparison of 80 to 90 genomes), the exact accessory genome (p-value < 0.008 in all cases), and the soft accessory genome (p-value< 0.02 in all cases). As was expected, orthogroups classified as exact and soft accessory genomes increased with the number of genomes included, ranging from 3360 to more than 11000 orthogroups.

**Figure 4.**
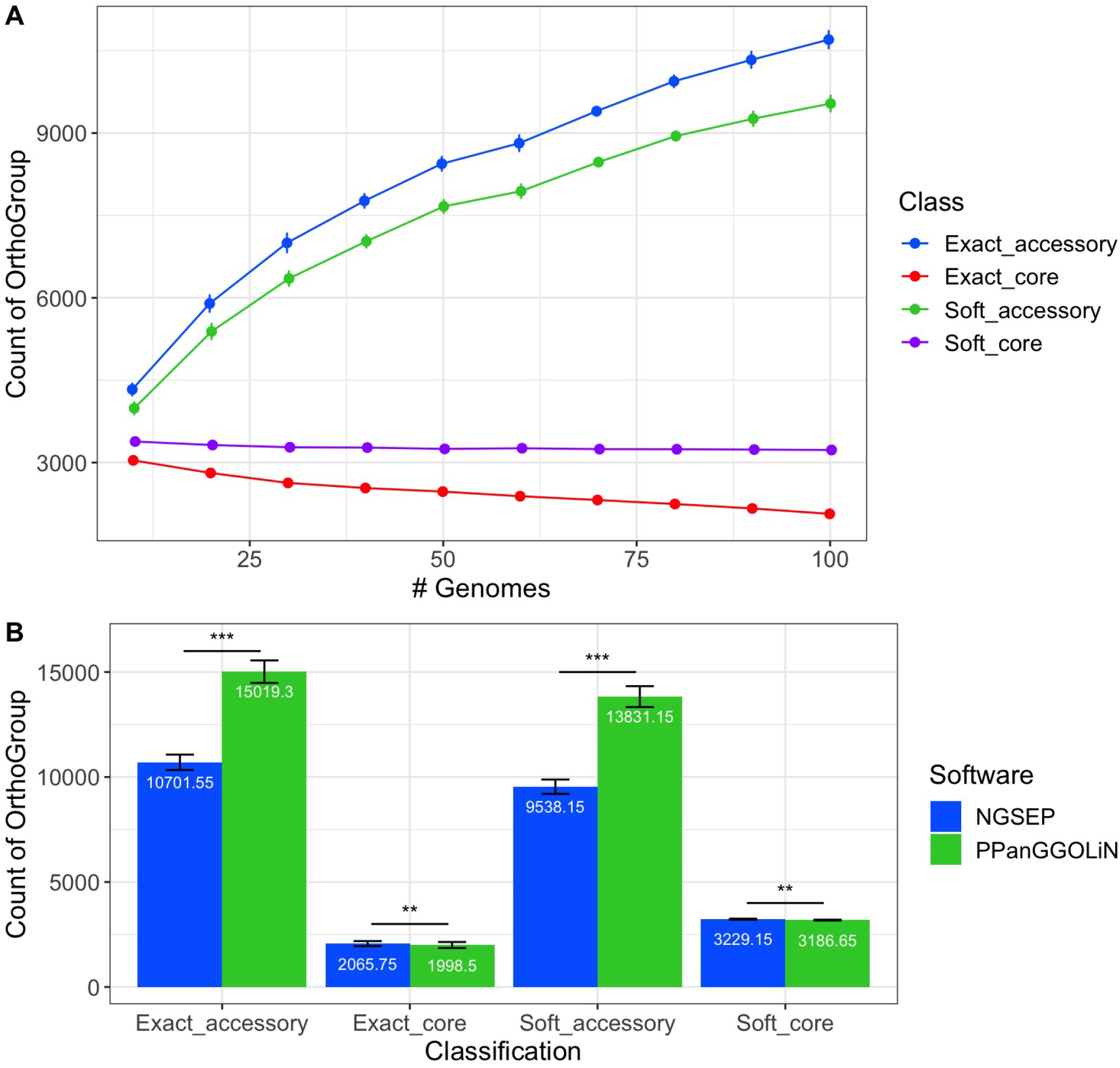
A)*Escherichia coli* pangenome reconstruction using 10 to 100 genomes from GenBank. Error bars show 95% confidence intervals. B) *Escherichia coli* pangenome reconstruction using NGSEP and PPanGGOLiN, and 100 genomes from GenBank. Error bars show standard deviation, ** p-value < 1e-5, *** p-value<1e-15

To compare our results with an open source tool, we performed the pangenome reconstruction using NGSEP and PPanGGOLiN on the subsets of 100 genome assemblies. The amount of orthogroups assigned to each partition are shown in Figure 4b. Due to the small 95% confidence intervals, the results showed a significant difference among the partitions in all cases. The significance values for all comparisons are available in Supplementary Table S3. Additionally, comparing the NGSEP soft-core genome with the persistent genome from PPanGGOLiN, the results also showed a significant difference on the mean (p-value 9.24e-10). NGSEP classifies about 30 more families as soft-core than PPanGGOLiN, and about 150 less families compared to the persistent classification. In contrast, PPanGGOLiN classifies about 5,000 more families as soft accessory, compared to NGSEP. The latter difference is caused by the algorithm for identification of orthologs and clustering method. In the case of PPanGGOLiN, it uses the searching algorithm from MMseqs, which includes a k-mer match step followed by a vectorized ungapped alignment and a gapped alignment (Steinegger and Söding 2017). After alignment, clustering is performed using a greedy set-cover algorithm (Hauser et al. 2016). These steps are previous steps for the classification algorithm. In the NGSEP implementation, the clustering is based on k-mer sharing and it can group distant gene families as was shown previously.

To understand the differences among classifications, the results from one dataset of 100 genomes were compared. A total of 9378 orthogroup clusters were common between NGSEP and PPanGGOLiN, 131 groups were unique in the PPanGGOLiN results, and 282 groups were only clustered with NGSEP. Additionally, 114 NGSEP groups were contained among PPanGGOLiN clusters, and 6942 PPanGGOLiN groups were included in the NGSEP clusters. Comparing the common orthogroups (See Supplementary Table S4), 2689 were classified as soft core by NGSEP; 2644 of them belong to the persistent partition in PPanGGOLiN, and 45 to the shell partition. Therefore, 6689 orthogroups were classified as soft accessory by NGSEP, only two of them belong to the persistent partition. These differences corresponded to orthogroups with frequencies between 89% and 94% that PPanGGOLiN reclassified according to the neighborhood information. We can conclude that NGSEP and PPanGGOLiN classifications had similar performances and the majority of differences are related to the clustering process.

#### Comparison of 16 mammals complete genomes

To assess if the algorithms implemented in NGSEP are able to perform ortholog identification and genomes alignment for large mammalian genomes, we aligned the assembled genomes of 16 mammal species, including 8 primates (5 hominids and 3 old world monkeys), 5 glires, and 3 outgroup mammal species (*L. africana, D. novemcinctus, and B. taurus*). Figure 5A, shows a literature-based reconstruction of the species tree. The 16 species have estimated divergence times ranging from 2.8 to 99 million years. The armadillo, elephant, and cattle species diverged from the other mammals more than 85 million years ago (Foley et al. 2016). The glires and the primate species separated 75 millions ago (dosReis et al. 2012, Foley et al. 2016). Overall, the glires have older estimated divergence times ranging from 73.5 to 15 million years ago. The divergence times of the analyzed primates are more recent, ranging from 29 to 2.8 million years ago. The group of hominids, to which *Homo sapiens* species belongs, diverged from other primates 29 years ago (Pozzi et al. 2014, Kumar et al. 2017). Humans separated from chimpanzees, the closest living evolutionary relatives, about 6.7 million years ago (Besenbacher et al. 2019, Hedges et al. 2015). The two chimpanzee species, *P. troglodytes* and *P. paniscus*, separated 2.8 million years ago (Schrago et al. 2012, Kumar et al. 2017).

**Figure 5.**
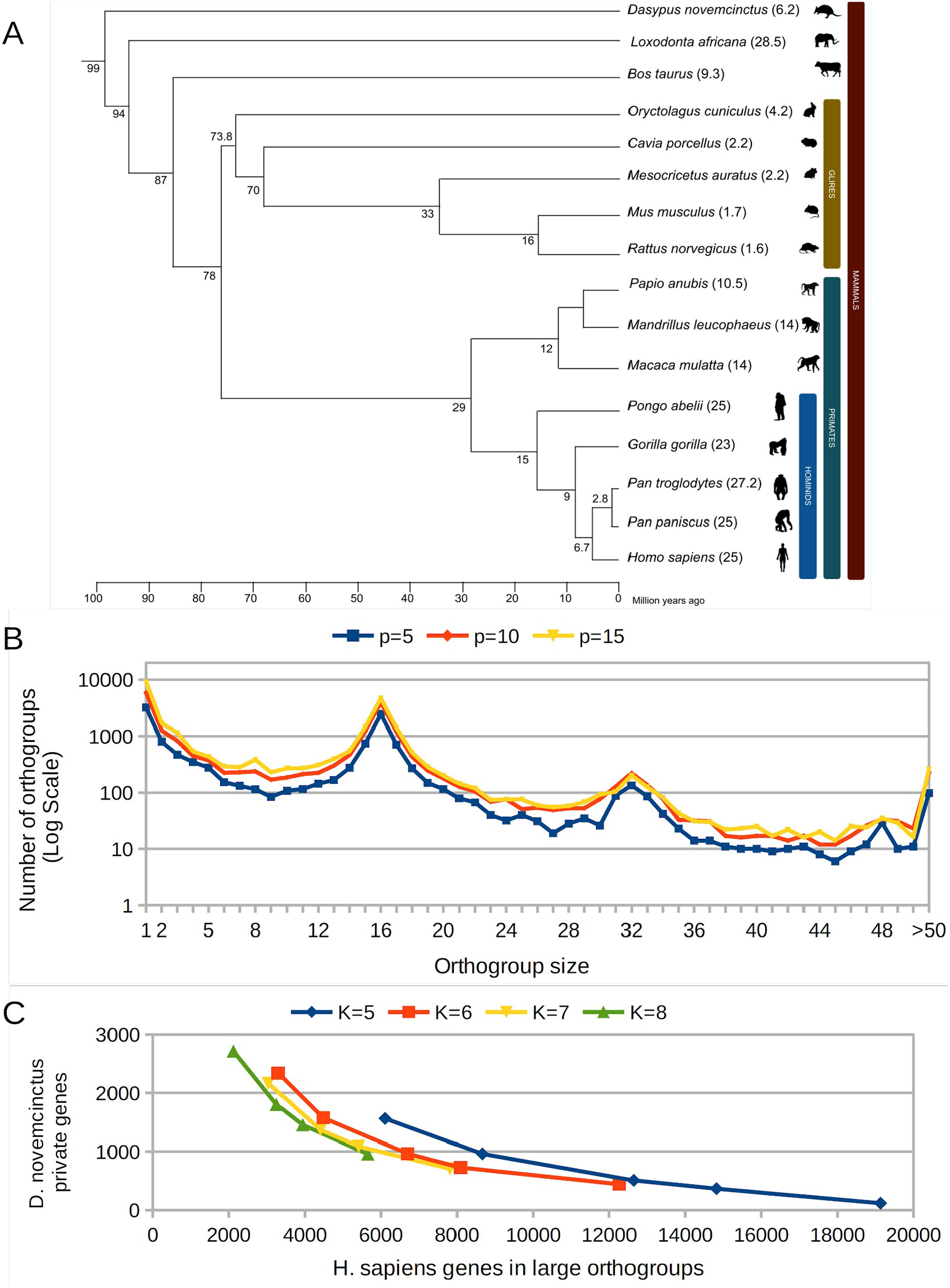
A Phylogenetic tree of analyzed Mammalian genomes. B Histogram of number of homolog clusters discriminated by cluster size for experiments with k=6 and varying the percentage of shared k-mers from 5% to 15%. C. Number of H. sapiens genes in large (size>100) orthogroups compared to the number of D. novemcinctus genes not included in an orthogroup. Each line is a series of experiments for a fixed k-mer length and varying the percentage of shared k-mers (5% to 20% for k=5 and k=6 and 5% to 15% for k=7 and k=8)

Taking into account the results of the benchmark experiments, the aligner was tested varying the k-mer length from 5 to 8, and varying the minimum percentage of shared k-mers from 5% to 20%. The distribution of cluster sizes consistently has a peak of up to 5000 orthogroups at the expected value of 16 for all experiments (Figure 5B). The distribution also shows two smaller peaks with around 300 and 200 orthogroups of size 8 and 32 respectively. The peak at size 8 corresponds to orthogroups specific to the 8 primate species. The peak at size 32 contains genes that are duplicated throughout the phylogeny.

The main difference among the analyses performed varying percentages is that the number of orthogroups at or close to all peaks increases as the percentage of shared k-mers increases (Figure 5B). The reason for this outcome is that at least one single large cluster (size larger than 100) is formed in all experiments. This cluster that merges several probably independent gene families is produced by running the connected components algorithm on a graph with probably false positive homolog relationships. For k=6, the number of human genes within large clusters decreases from 12,268 genes to 3,296 genes as the percentage increases from 5% to 20% (Figure 5C). This abnormally high number of presumably genes with paralogs is not obtained when the analysis is performed solely on the human genome. The largest cluster obtained in this case is 1,613 using a minimum 5%, and this cluster is not retained if the percentage is increased to 10%. This indicates that false positive orthologs can generate false paralog relationships that are not observed in a single genome analysis.

Conversely, increasing the percentage for the analysis including the 16 species increases the number of genes that can not be included in an orthogroup. For *D. novemcinctus* and k=6, the number of presumably private genes increases from 443 to 2,335 genes as the percentage increases from 5% to 20%. This number of private genes is not obtained for *H. sapiens* because the analysis includes species such as *P. troglodytes*. For *H. sapiens*, the number of presumably private genes remains below 30, even at p=15%. Figure 5C shows a tradeoff between the number of *H. sapiens* predicted paralogs (as an estimate of false positive relationships) against the number of predicted private genes for *D. novemcinctus* (as an estimate of false negative relationships). Results for different k-mer lengths are plotted as lines generated varying the percentage of shared k-mers. Although increasing the k-mer length seems to improve the tradeoff in this comparison, the difference between k=6 and k=8 is small compared to that between k=5 and k=6.

## DISCUSSION

Comparative analysis of large numbers of accurate genome assemblies between and even within species requires a new generation of scalable software tools to reveal significant results on evolutionary and functional genomics. In this work, we aim to contribute to this challenge providing innovative algorithms and functionalities for common tasks in comparative genomics such as identification of homology relationships across genes and whole-genome alignment. Our work was inspired by recent manuscripts describing genome comparisons (Zhang et al. 2016, Stein et al. 2018, Moghaddam et al. 2021), which needed to combine different tools to perform ortholog identification and genome alignments. Our goal in this work was to provide a tool able to call orthologs and compare genome assemblies in the length range of the human genome with good usability, high accuracy and low computational resources.

Considering the described needs and that even current tools such as Orthofinder rely on BLAST/DIAMOND searches to predict raw homology relationships, we started by implementing a novel approach for an efficient identification of homology among protein sequences. The indexing and comparison of amino acid k-mers represented a big improvement in computational efficiency, compared to BLAST searches, with an almost negligible penalty on accuracy. K-mer co-occurrence methods, also known as alignment free methods (Chan and Ragan 2013) have been proposed as an efficient proxy for genome alignment scores for detecting homology in large proteomes of bacteria, fungi, plants and Metazoa (Anari et al. 2018). We also contributed with integrated implementations of the Markov Clustering algorithm for identification of gene families, and the HalSynteny algorithm for identification of synteny blocks and genome alignment. These new implementations facilitate the installation and execution of the entire analysis pipeline as a single step. Moreover, the Markov Clustering is only executed on connected components, which significantly increases the computational efficiency of the entire solution compared to pipelines integrating OrthoMCL as an external software tool.

Benchmark experiments comparing the NGSEP genomes aligner with other software tools for identification of orthogroups indicate that NGSEP has competitive accuracy compared to OrthoFinder (Emms and Kelly 2019), better accuracy compared to SonicParanoid (Cosentino and Iwasaki 2019), and the best computational efficiency observed. We acknowledge that part of the larger computational execution time required by Orthofinder2 is explained by the extra processing needed to build both a species tree and gene trees for the orthogroups. However, our experiments with the subset of mammalian gene families from Orthobench suggest that the potential improvement in accuracy of orthogroups that could be achieved by these extra processes requires a wide representation of the phylogeny of the species under study. Conversely, the genomes aligner of NGSEP performs an efficient ortholog clustering for collections of genome assemblies separated by relatively short evolutionary times. We believe that this is a very common use case for genome assembly projects currently under development. Our experiments indicate that the algorithm implemented in NGSEP is robust to changes in the representation of the taxonomy under study. The difference between the results obtained with the Orthobench dataset and the complete mammal genomes trying different parameter suggests that the orthobench dataset needs to be improved to achieve a better assessment of potential false positive relationships, and that additional benchmark strategies or datasets should be developed to perform a comprehensive assessment of the accuracy of tools for orthogroup identification.

We identified a wide range of perspectives and future software development works for comparative genomics. As improvements to the current functionalities, we believe that alternative clustering techniques could be explored to improve the efficiency of orthogroup identification. Although our solution integrates identification of orthogroups and synteny blocks, the synteny blocks could be further used to increase accuracy of orthogroup identification. Regarding new functionalities and taking into account the path followed in the development of Orthofinder, we are currently exploring alternatives to build species and gene trees, exploring the additional synteny information that we are obtaining from contiguous genome assemblies. Moreover, in this work we provide a basis for construction of gene-based pangenomes within NGSEP. We are currently exploring more sophisticated alternatives to classify gene families and increase the accuracy of this process. Most recent works are focused on building sequence-based instead of gene-based pangenome graphs (Minkin and Medvedev 2020). We hypothesize that a gene-based pangenome could be used as an initial step to increase efficiency in the construction of sequence-based pangenome graphs.

We have used different earlier versions of the described functionalities to identify orthologs and large genome rearrangements between common bean and Lima bean (Garcia et al. 2021) and to identify orthologs between sugar cane and different close species (Trujillo et al. 2021). We expect that the new functionalities for comparative genomes presented here would be used by a large number of research groups currently building and comparing genome assemblies in a wide variety of species.

## METHODS

### Setting parameters for Orthogroup Inference

The Orthobench dataset (Emms and Kelly 2020) was used to determine the best algorithm parameters and compare our software against other solutions. Briefly, this database contains 70 reference orthogroups (RefOGs) manually curated by experts that span the Bilateria organisms and covers a range of different challenges for orthogroup inference (All information is available at https://github.com/davidemms/Open_Orthobench). As a first approach, the percentage of 3 to 9-mers shared by each RefOG was calculated, allowing us to identify a suitable k-mer percentage to identify orthologs. However, this approximation is only useful for finding the parameters that end in a good recall but not in a good precision. Therefore, all 70 RefOGs were separated by species, resulting in 12 Bilateria proteome input files. These files were processed to identify orthogroups and to establish the recall, precision, f-score, and execution time of the algorithm. K-mer lengths from 5 to 9-mer, and percentages of shared k-mers from 1 to 10% were tested. The best parameters were set as default parameters of NGSEP to compare with other tools.

To compare more closely related species, the Bilateria dataset was filtered to include only mammal species. This dataset was also tested using k-mer lengths from 5 to 9-mer, and percentages of shared k-mers from 1 to 10%. Finally, resulting clusters were compared to the database clusters to calculate recall, precision, and f-score.

### Benchmarking orthologs

To compare our algorithm to commonly used software tools, we performed the orthologs identification using NGSEP, OrthoFinder and SonicParanoid. These tools were selected because of their top rankings in the Orthobench benchmarking service. Our algorithm was tested with the Markov Clustering step using 5-mer and 5% of shared k-mers. OrthoFinder was tested using the first and second versions, with BLAST and DIAMOND pairwise comparisons. SonicParanoid was tested with the default parameters, as well as the most-sensitive configuration. Comparisons were made with the complete dataset (70 RefOGs, 12 Bilateria species), and the filtered dataset (mammals). In all cases, recall, precision and f-score were calculated versus the curated clustering.

### Comparison of bacterial gene families

To test our algorithm over complete genomes, we performed a clustering and pangenome reconstruction using bacterial datasets. A total of 15000 *Escherichia coli* genomes were retrieved from GenBank; these genomes accomplished the quality criteria previously defined by Gautreau et al. (Gautreau et al. 2020). The complete list of genomes is available in the Supplementary Table S5. Due to the sensitivity of the orthologs identification and pangenome reconstruction to the number of genomes included, 10 to 100 genome batches were randomly subsampled, in increments of 10 genomes, for a total of 20 groups for each batch. Then, GFF3 annotation files and FASTA files were used as input for the NGSEP Genomes Aligner fixing the k-mer size to 6 and the percentage of shared k-mers to 5%, resulting in a set of text files composed by the clusters (orthogroups), presence/absence matrix and orthogroups frequencies among the genomes. Based on frequencies, the exact/soft classification into core and accessory genome was obtained, using default parameters (90% threshold for the soft classification). A t-test was performed to compare the number of orthogroups assigned to core and accessory genomes among subsampled groups.

To compare our results to open-source software, we used the 100 genomes batches to run PPanGGoLiN v1.2.63 (Gautreau et al. 2020). The software was configured to run with the – annot option and the GenBank files as input for the algorithm. The amount of gene families classified as exact/soft core and accessory, were calculated and compared to the amounts of NGSEP. A t-test was performed to compare the number of orthogroups assigned to each partition between the software tools.

### Comparison of 16 mammals complete genomes

We retrieved the genomes and their annotation from 16 phylogenetically diverse mammals from ENSEMBL, including 5 hominids: human (*Homo sapiens)*, bonobo (*Pan paniscus)*, chimpanzee (*Pan troglodytes)*, gorila (*Gorilla gorilla)*, and orangutan (*Pongo abelii)*; three old world monkeys: rhesus macaque (*Macaca mulatta)*, drill (*Mandrillus leucophaeus)*, and olive baboon *(Papio anubis*); 5 glires: brown rat (*Rattus norvegicus)*, house mouse (*Mus musculus)*, guinea pig (*Cavia porcellus)*, golden hamster (*Mesocricetus auratus)*, and European rabbit *(Oryctolagus cuniculus*); 1 ungulate: cattle (*Bos taurus*); 1 pen-ungulate: African elephant (*Loxodonta africana*); and 1 Xenarthra species: nine-banded armadillo (*Dasypus novemcinctus*). The complete list of genomes and downloaded links is available in the Supplementary Table S6. The species tree (Figure 5A) was built combining results from different sources using TimeTree (Kumar et al. 2017).

## COMPETING INTERESTS STATEMENT

The authors declare that there are no competing interests related to the results presented in this manuscript

## ACKNOWLEDGEMENTS

This work has been supported by the Colombian research fund “ PATRIMONIO AUTÓNOMO FONDO NACIONAL DE FINANCIAMIENTO PARA LA CIENCIA, LA TECNOLOGÍA Y LA INNOVACIÓN FRANCISCO JOSÉ DE CALDAS” through the grant with contract number 80740-441-2020, awarded to JD. This work was also supported by internal funds of Universidad de los Andes through the FAPA initiative led by the Vice-presidency of Research and Knowledge Creation. We also wish to acknowledge the support of the DSIT high performance computing unit at Universidad de los Andes to conduct part of the bioinformatic analysis presented in the manuscript.

## Notes

### Competing Interest Statement

The authors have declared no competing interest.

